# Spatial autocorrelation of species diversity and distributions in time and across spatial scales

**DOI:** 10.1101/2025.06.13.659470

**Authors:** Carmen D. Soria, Gabriel R. Ortega, Friederike J.R. Wolke, Vojtech Bartak, Katerina Tschernosterova, Vladimir Bejcek, Sergi Herrando, Ivan Mikulas, Karel Stastny, Mutsuyuki Ueta, Petr Vorisek, Petr Keil

## Abstract

**Aim:** Spatial autocorrelation (SAC), also known as aggregation, is a notable property of species distributions and diversity; it reflects species niche and dispersal, has conservation significance, and affects ecological models. Yet, we know little about spatial and temporal patterns of SAC in empirical data. Here, we assess SAC in both observed species distributions and species richness, quantifying its magnitude and prevalence over large extents and across spatial resolutions. We also assess its dynamics over the past 50 years.

**Location:** Czechia, Europe, New York State, Japan

**Time period:** 1972 - 2017

**Major taxa studied:** Birds

**Methods:** We analyzed four temporally replicated gridded bird atlases, each aggregated to multiple grain sizes. To measure SAC in species distributions, we used the Join count statistic (JC) and its deviation from the expectation under a random distribution. We assessed temporal changes in JC and their relationship with changes in occupancy, given their close association. We used Moran’s I to measure SAC in species richness.

**Results:** Both species distributions and diversity were positively autocorrelated across all regions, periods, and grains, and the magnitude of autocorrelation mostly decreased with increasing grain. We found that the temporal change of JC varied across species and regions, with zero average trends in Moran’s I, JC, and occupancy. However, when JC and occupancy were considered jointly, we found systematic temporal shifts: contracting species became more aggregated (compact) while expanding species became more fragmented (disjoint).

**Main conclusions:** Stronger SAC at finer grains suggests greater predictability of diversity and distributions at these scales. Despite zero average change in occupancy or SAC, their coupled shifts highlight the importance of considering both jointly. We found long-distance dispersal (rather than advancing edge) and vulnerability of isolated populations to extinction as the major drivers of range dynamics in temperate birds.

## Introduction

Macroecology has historically focused on static patterns of biodiversity and species distributions (Brown, 1995; Rosenzweig, 1995), but human-driven global change has likely been altering these patterns (IPBES, 2019). Thus, research on temporal changes in species diversity and distributions has been increasing, including studies on temporal changes in single-species population abundance (Rosenberg *et al*., 2019; Ledger *et al*., 2023), occupancy (Warren *et al*., 2001; Klinkovská *et al*., 2024), and range position (Thomas & Lennon, 1999; Chen *et al*., 2011). Multi-species studies have predominantly examined changes in species richness (Vellend *et al*., 2013; Dornelas *et al*., 2014; Blowes *et al*., 2019), assemblage composition (Jones *et al*., 2020), interspecific associations (Calatayud *et al*., 2019; Keil *et al*., 2021), and turnover (Blowes *et al*., 2024). However, a comprehensive large-scale assessment of spatial and temporal variation in the *spatial aggregation* and *fragmentation* of species distributions and diversity, which can be measured by their spatial autocorrelation (SAC, Box 1), has been missing.

### Spatial autocorrelation as a summary statistic

SAC, which indicates the degree of spatial dependence among values of a variable, is often considered a nuisance in ecological modeling (Lennon, 2000; Dormann, 2007). Autocorrelated observations are typically viewed as a form of pseudo-replication, reducing the degrees of freedom (Fortin & Dale, 2005), violating the assumption of independent residuals, biasing parameter estimates, and increasing Type I error rates (Dormann *et al*., 2007). An alternative view is that SAC is a useful descriptive *summary statistic* (*sensu* Wiegand & Moloney, 2013) of ecological systems, which can provide additional information to metrics such as species occupancy or richness, potentially informing on the processes shaping biodiversity and species distributions (Dormann *et al*., 2007; Hawkins, 2012). The SAC of species distributions can also be viewed as an indicator of range fragmentation (or its opposite, i.e., aggregation). High SAC values indicate less range fragmentation and higher distribution integrity, whereas low SAC values indicate increased fragmentation and reduced connectivity between populations.

Examples of ecological studies that summarize species patterns of SAC include Koenig, 2001; Cocu *et al*., 2005; Bosiacka *et al*., 2012; Chevalier *et al*., 2021 and Moctezuma, 2021. In terms of ecological inference, SAC has been used to identify spatial discontinuities between communities and the spatial extent of inter-species associations (Hui, 2009; Bonada *et al*., 2012), to disentangle exogenous and endogenous drivers of SAC (Bonada *et al*., 2012; Mielke *et al*., 2020), to test whether SAC decay in species occupancy matches that of macroclimatic variables (Rich & Currie, 2018), to describe spatiotemporal distributions of invasive species (Barney *et al*., 2008; Wang *et al*., 2011), or to describe the spatial extent of SAC (Koenig, 2001; Chevalier *et al*., 2021; Niwa & Uno, 2023). Given SAC’s potential utility, it is surprising that it has neither been systematically assessed over large geographic extents of empirical data and across many species, nor in the context of biodiversity change.

### Spatial autocorrelation and species occupancy in time

A fundamental property of every species is its *occupancy*, i.e., the number (or proportion) of sites it occupies in geographic space. Occupancy is a key facet of species rarity (Crisfield *et al*., 2024), range size (Orme *et al*., 2006), and extinction risk (IUCN Standards and Petitions Committee, 2024). Thus, assessing temporal change of occupancy is highly relevant in large-scale biodiversity assessments (Warren *et al*., 2001; Jetz *et al*., 2019; Klinkovská *et al*., 2024).

Occupancy is closely linked to SAC: in a limited space, the higher the occupancy, the higher the chance occupied locations will be near each other, even if distributed randomly. Consequently, high occupancy can lead to high measured SAC for reasons that are not ecological but a geometrical inevitability. To account for this, studies of the SAC of species distributions should report both occupancy and SAC together (Fig. 1a; Niwa & Uno, 2023) or use Z-scores, which quantify the deviation of the observed SAC from its expected value under randomly distributed occupied sites (Fig. 1a; Lee, 2003; Wang *et al*., 2011; Niwa & Uno, 2023). However, since the relationship between occupancy and SAC is not deterministic — a single occupancy value can have multiple SAC values — observed changes in SAC can deviate from the expected based on occupancy change alone, resulting in *overaggregation*, where SAC increases more or decreases less than expected, or in *overdispersion*, where SAC decreases more or increases less than expected for the observed change in occupancy (Fig. 1b). Changes in SAC can also be fully independent of changes in occupancy, with species distributions becoming more spatially aggregated or fragmented even when occupancy remains stable (Fig. 1a). Therefore, solely focusing on the total area a species occupies following a disturbance may potentially obscure the existence of severe range fragmentation.

**Figure 1.**
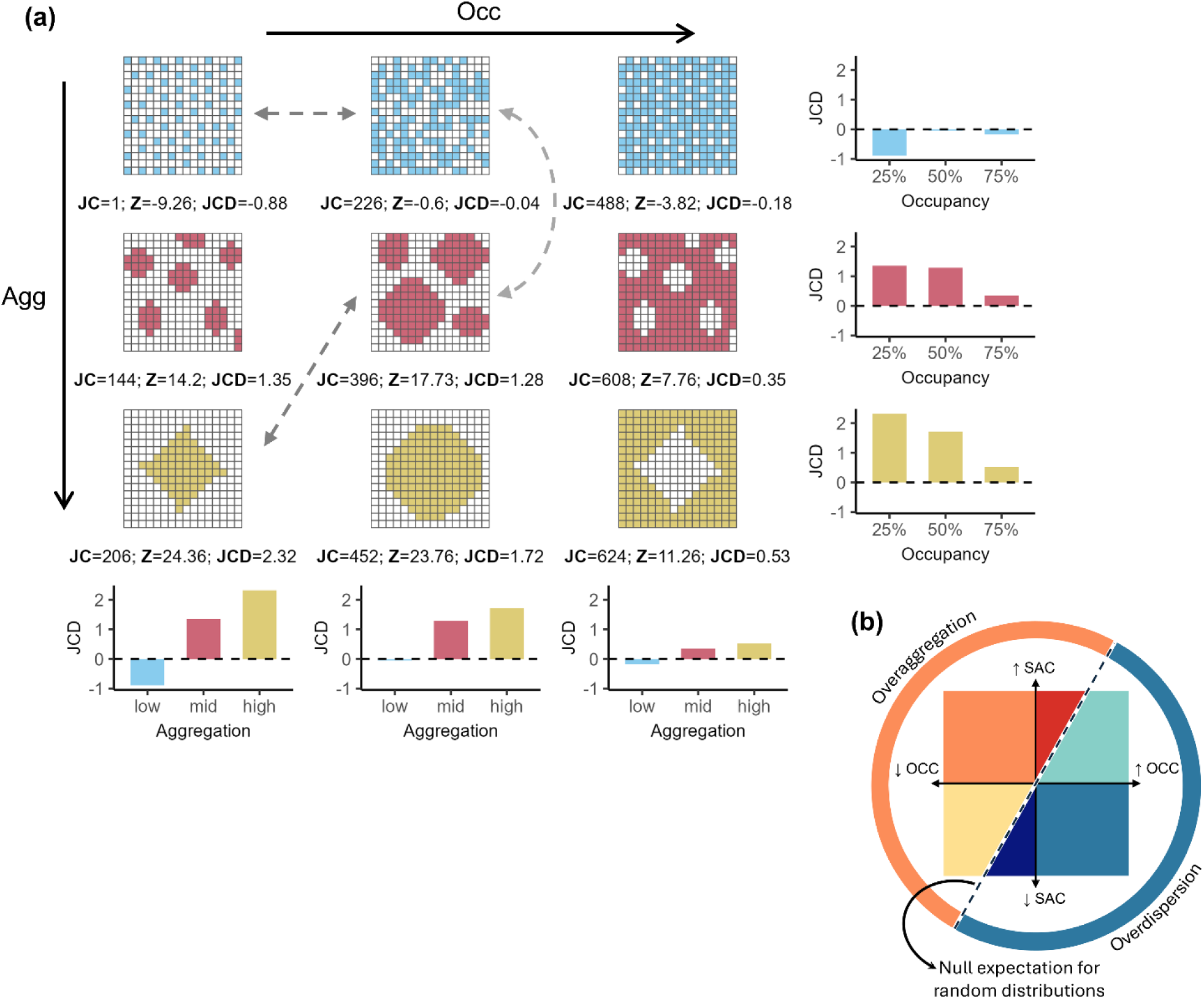
Relationship between temporal changes in SAC and occupancy. Panel (a) shows the spatial distribution of a virtual species, where colored cells are presences and white cells are absences. Occupancy (columns: 25%, 50%, 75%) and aggregation levels (rows: low in blue, medium in red, high in yellow) vary independently. Species distributions may shift along the occupancy axis (horizontal), aggregation axis (vertical), or both, as indicated by the grey dashed arrows. Spatial autocorrelation (SAC) is quantified using three metrics (see Methods): Join count (JC), measuring the number of adjacent presences; Join count Z-score (Z), assessing whether presences are significantly aggregated (Z > 1.96) or dispersed (Z < -1.96) at a 95% confidence level; and average Join count difference (JCD), a metric of SAC that accounts for occupancy and number of cells. Panel (b) decomposes the expected relationship between changes in occupancy (OCC; x-axis) and spatial autocorrelation (SAC; y-axis) in species distributions. The black dashed line is the expected SAC change for a given occupancy change. Deviations from this line indicate a higher increase or lower decrease in SAC (overaggregation; orange semi-circle) or a lower increase or higher decrease in SAC (overdispersion; blue semi-circle) than expected.

Changes in individual species distributions can also affect the spatial structure of diversity. Similar to species distributions, the SAC of richness can vary independently of total richness, reflecting shifts in its spatial arrangement. If areas of high species richness become more clustered (i.e., spatially structured), SAC will increase. On the other hand, if species richness becomes more evenly or randomly distributed, SAC will decrease. We, therefore, argue for empirically assessing temporal trends in the SAC of both species’ distributions and richness.

### Spatial autocorrelation and spatial scale

Another critical gap in our understanding of SAC is its relationship with spatial scale, particularly spatial grain—the average size of the elementary sampling unit (Legendre & Legendre, 2012). Species occupancy has a non-linear relationship with grain size. As grain size increases, neighboring cells merge, resulting in an increase in the proportion of occupied cells that eventually levels off, a phenomenon named the scaling pattern of occupancy (Kunin, 1998; Hartley & Kunin, 2003; Hui *et al*., 2006). Similar to occupancy, SAC varies with spatial grain.

Theoretical models show that the SAC of species distributions decreases with increasing grain size, while the overall spatial structure (i.e., aggregated or dispersed) remains unchanged (Hui, 2009). The factors driving SAC are also grain dependent: at fine-grains, SAC is primarily driven by endogenous processes, though microhabitats may also contribute; at coarser grains, broad-scale environmental (i.e., exogenous) factors become more prominent, with long-distance dispersal events potentially playing a role too. Using a virtual species approach, Fig. 2 illustrates that SAC can increase or decrease with grain size, depending on the spatial pattern of the fine-scale distribution. It remains unclear which of these SAC patterns (i.e., increasing or decreasing with grain) is most prevalent in nature. The total number of available sites should also be considered, as areas with more sites will have a lower expected SAC under a random distribution. Namely, clustering is more noticeable when there are many sites available than few, and as grain size increases, the number of sites decreases, influencing whether observations are significantly spatially autocorrelated.

**Figure 2.**
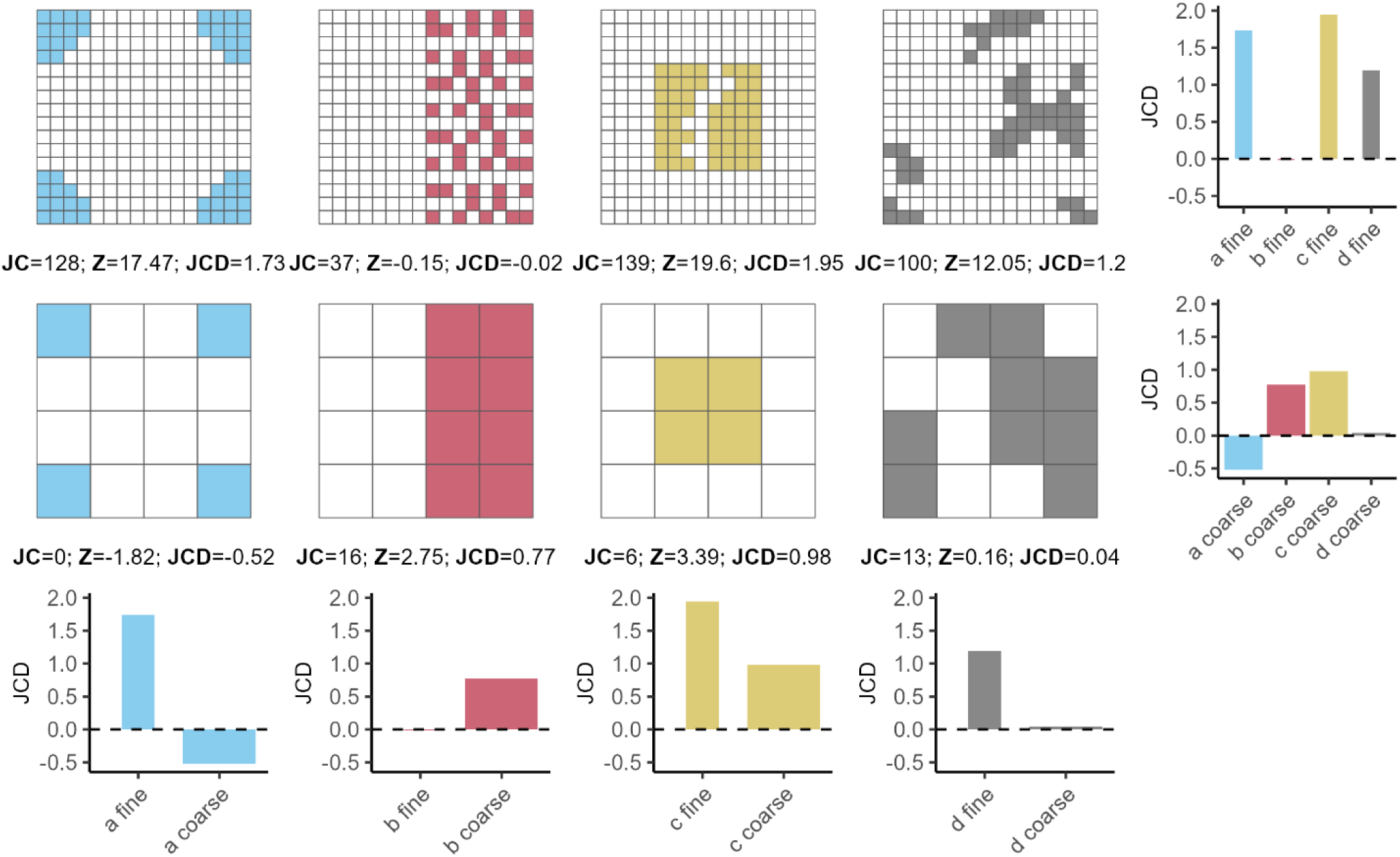
Spatial autocorrelation (SAC) across spatial grains. The first row presents four hypothetical fine-scale species distributions, each with an occupancy of 20.31% (52 presences). The second row shows the coarse-scale versions of each scenario. We quantified the SAC of species’ presences for each scenario using three metrics (see Methods): Join count (JC), measuring the number of adjacent presences; Z-score (Z), assessing whether presences are significantly aggregated (Z > 1.96) or dispersed (Z < -1.96) at a 95% confidence level; and average Join count difference (JCD), a variation of JC that accounts for the effects of occupancy and number of cells. Barplots show JCD values for each scenario grouped by grain size and distribution structure.

For species richness, SAC is similarly expected to vary with grain size, as the spatial pattern of richness results from overlaying individual distributions, each with its own grain-dependent SAC. As grain size increases, total richness typically rises as larger areas can host more species. This can result in an increase in SAC if smooth gradients emerge at that grain size. Conversely, SAC declines if richness becomes more uniform or random and less spatially structured. To the best of our knowledge, no studies have analyzed the effect of grain size on the SAC of species distributions and diversity across large geographical extents.

### Objectives

Here, we aim to explore SAC as a facet of biodiversity (i.e., as a summary statistic) that can be analyzed alongside properties such as species richness, occupancy, or beta diversity within the context of global change. We ask the following questions: (1) Are empirical species richness and distributions spatially autocorrelated, independent of occupancy? (2) How has empirical SAC changed over time, separate from changes in occupancy? (3) How does empirical SAC vary across spatial grain sizes, independent of occupancy? (4) What is the relationship between changes in occupancy and corresponding changes in SAC? To achieve this, we assess patterns of SAC in several large avian datasets with continuous spatial coverage and temporal replications, specifically four Breeding Bird Atlases (BBAs).

##### Box 1: What is spatial autocorrelation?

Species diversity and distributions are not random but spatially aggregated or fragmented, a phenomenon known as spatial autocorrelation (SAC; Legendre, 1993). Spatial autocorrelation measures the similarity of observations as a function of spatial distance. It can be *positive*, where geographically close observations are more similar than those further apart, or *negative*, where nearby observations are more dissimilar (Legendre, 1993). Positive SAC indicates clusters of species’ presences or richness values, while negative SAC indicates spatial dispersion. Based on the underlying processes, the SAC of species distributions can be classified into two main types: *exogenous* and *endogenous* (Dormann, 2007). Exogenous SAC arises from autocorrelated external environmental drivers such as climate, land use, and topography (Fortin & Dale, 2005; Dormann, 2007). In contrast, endogenous SAC emerges from processes intrinsic to species, such as population dynamics, competition, dispersal, or localized movement (Fortin & Dale, 2005; Dormann, 2007).

Several metrics have been developed to measure SAC, each addressing different aspects of spatial dependency. *Global metrics*, such as Moran’s I (Moran, 1950) and Geary’s c (Geary, 1954) for continuous or count variables, or the Join count statistic (Sokal & Oden, 1978) for binary or categorical variables, summarize overall spatial patterns in a single number, indicating whether similar values aggregate or disperse across the study area. *Local metrics*, such as Local Indicators of Spatial Association (LISA, Anselin, 1995) and Local Indicators for Categorical Data (LICD, Anselin & Li, 2019), identify locations of high or low SAC. Additionally, correlograms and semi-variograms visualize SAC by plotting similarity (correlograms) or variance (semi-variance) against geographic distance (Dormann *et al*., 2007).

## Materials and Methods

### Species distribution data

Bird distribution data were obtained from four Breeding Bird Atlases (BBAs), which use standardized breeding-season surveys across grid cells covering geopolitical regions such as cities, provinces, or countries. Surveys are coordinated by experts and carried out by volunteers and professionals, following an established protocol specific to each atlas. Each sampling period spans multiple years with repeated visits to most cells. Some regions have temporal replications of surveys (atlas replications), allowing for more reliable comparisons of species occurrences over time than nonstandardized surveys. While true absences are hard to confirm, BBAs offer one of the best large-scale approximations of presence–absence data (Pototsky & Cresswell, 2023).

We utilized BBAs from Czechia, Europe, New York State (hereafter New York), and Japan. These atlases cover diverse geographical areas in the Northern Hemisphere, species pools, time periods, survey durations, spatial extents, and grain sizes (Table 1). Each atlas has at least two replications, enabling temporal analyses of SAC. The first Czech BBA (1973 – 1977) was not considered due to grid incompatibility with subsequent periods. The third Japanese BBA (2016 – 2021) was also excluded due to a substantial increase in voluntary reports compared with previous atlas replications.

**Table 1.** Summary of the four BBAs used in the empirical assessment of patterns of spatial autocorrelation (SAC).

| Location | Replications | Periods | Grain size | Area | References |
| --- | --- | --- | --- | --- | --- |
| Czechia | 3 | 1985 – 1989,<br>2001 – 2003,<br>2014 – 2017 | ~ 11x11 km<br>(10' long x 6' lat) | ~ 78,309 km<sup>2</sup> | (Šťastný et al., 1997, 2006, 2021) |
| Europe | 2 | 1972 – 1995,<br>2013 – 2017 | ~ 50x50 km | ~ 5,909,158 km<sup>2</sup> | (Hagemeijer & Blair, 1997; Keller et al., 2020) |
| New York State | 2 | 1980 – 1985,<br>2000 – 2005 | ~ 5x5 km | ~ 126,879 km<sup>2</sup> | (Andrle & Carrol, 1988; McGowan & Corwin, 2008) |
| Japan | 2 | 1974 – 1978,<br>1997 – 2002 | ~ 20x20 km | ~ 367,613 km<sup>2</sup> | (Biodiversity Center of Japan, 2004; Environment Agency of Japan, 1981) |

### Data processing

BBA data consist of species presence records associated with specific grid cells over defined atlas periods. We kept cells surveyed in all atlas periods and excluded species records flagged as uncertain by data curators (see Supporting Information). We aggregated grid cells by doubling the grid cell side at each coarsening step (i.e., 2x2, 4x4, 8x8, etc.), considering a coarser cell as surveyed if at least one of its constituent fine-grain cells had been surveyed. A species was considered present in the coarser cell if recorded in any of the smaller constituent cells. Additionally, we matched the taxonomy of each BBA to the HBW/BirdLife Taxonomic Checklist (2024, version 9.0).

Species occupancy and grid cell level richness were calculated for each atlas region, period, and grain size combination. Only grain sizes with at least 30 cells were considered for SAC analysis (Lee, 2003). The minimum and maximum grid cell resolutions analyzed were 11x11 and 44x44 km for Czechia, 50x50 and 800x800 km for Europe, 5x5 and 80x80 km for New York, and 20x20 and 160x160 km for Japan (Fig. 3).

**Figure 3.**
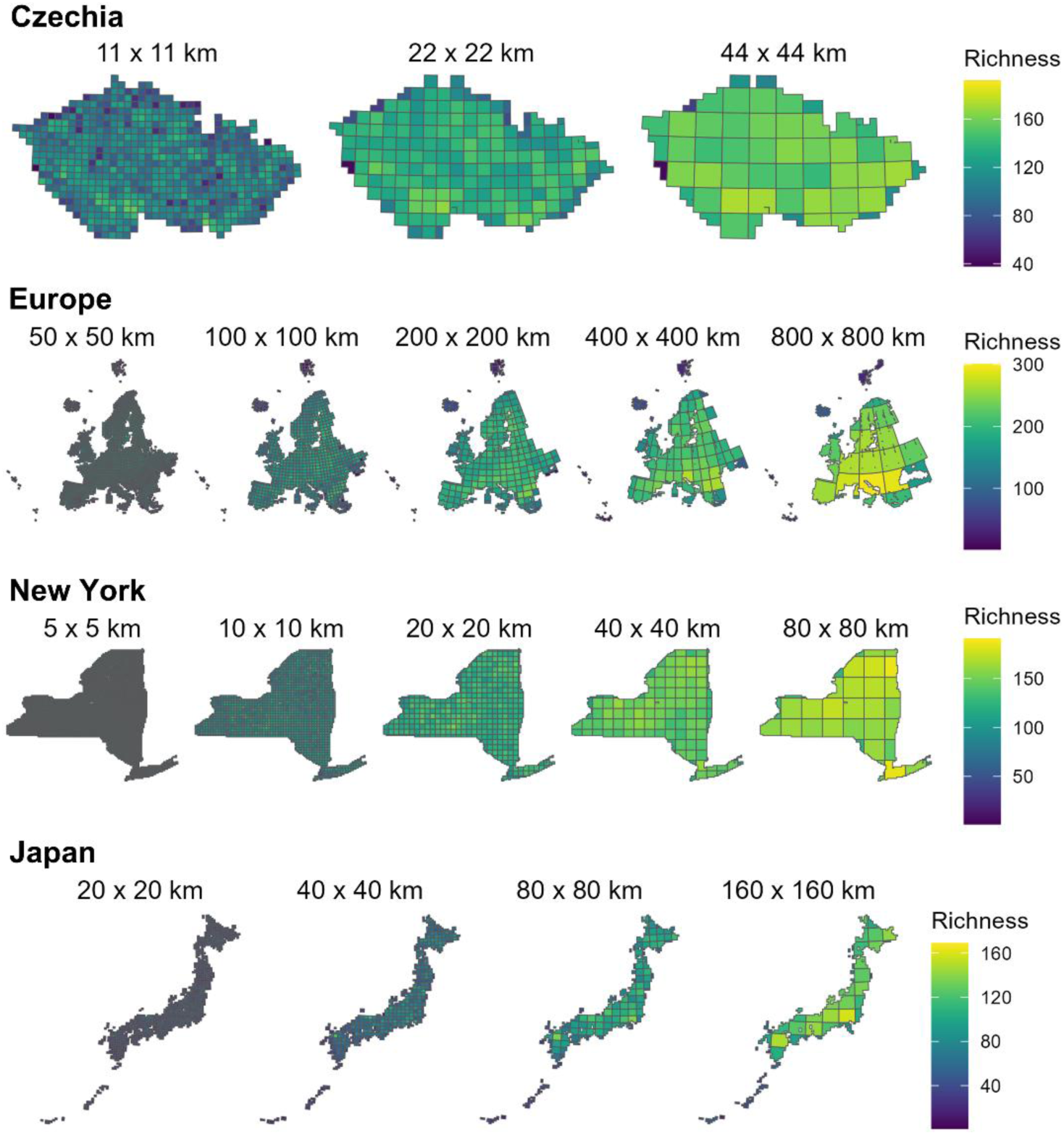
Species richness in the first atlas period for all atlas regions and grain sizes. Each panel displays species richness, represented by a color gradient, aggregated at progressively larger spatial resolutions. Richness increases from blue (low) to yellow (high). The area of each cell aggregation (i.e., grain size) is indicated on top of each map. All maps were projected using the Lambert Azimuthal Equal-Area (LAEA) projection, with central meridians and standard parallels specific to each area to minimize distortion.

### SAC in species diversity and distributions

We assessed the global SAC for species richness using Moran’s I and for species distributions using the Join count statistic (JC) across each combination of atlas region, replication, and grain size.

*Moran’s I* quantifies how similar values cluster together in space for continuous and count data, such as species richness. The relationship between locations is defined through a spatial weights matrix 𝑊, generally giving more weight to nearer locations, and is calculated as:

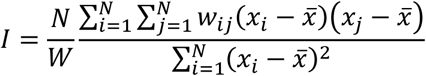

Where 𝑁 is the number of spatial units, 𝑥_𝑖_ and 𝑥_𝑗_ are the values of the observations at locations 𝑖 and 𝑗, <u>𝑥</u> is the mean of the variable of interest across all locations, 𝑤_𝑖𝑗_ is the spatial weight between locations 𝑖 and 𝑗, and 𝑊 is the sum of all spatial weights 𝑤_𝑖𝑗_. Moran’s I values range from -1 to +1, where negative values indicate dissimilarity between neighbors, positive values suggest an aggregation of similar values, and values near 0 reflect a random spatial pattern. We calculated Moran’s I values considering all immediately adjacent neighbors (i.e., first distance class queen contiguity) and using a row-standardized weighting scheme in which all neighbors sum to 1 (Bivand *et al*., 2013). Moran’s I significance was assessed by comparing the observed value to its distribution expected under the null hypothesis of no spatial autocorrelation.

*Join count statistic (JC)* measures SAC in binary or nominal variables, such as presence-absence data. When considering species presences, a “join” occurs when two neighboring sites are occupied (similarly, when considering absences, two neighboring unoccupied sites constitute a join). A high number of joins indicates clustering (aggregation) of presences and positive SAC. Mathematically:

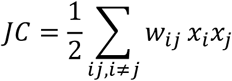

𝑥_𝑖_ and 𝑥_𝑗_ are the values of the observations at locations 𝑖 and 𝑗, and 𝑤_𝑖𝑗_ is the spatial weight between locations 𝑖 and 𝑗. It can only be calculated for distributions with both presences and absences. Similarly to Moran’s I, the significance of the JC is calculated by comparing the observed value to its expected value under the null hypothesis of no spatial autocorrelation. We calculated the number of observed joins considering first distance class queen contiguity with a binary weighting scheme in which all neighboring cells have the same weight (Bivand *et al*., 2013). The expected number of joins and their variance were calculated under a non-free sampling assumption, where the number of presences is fixed, but their spatial arrangement can vary. These expected values represent the number of joins we would expect if presences and absences were randomly distributed across the study area.

Because the JC is influenced by occupancy and the total number of available sites, we used the average Join count difference (hereafter JCD) instead of the raw JC, defined as:

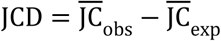

Where 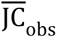 and 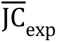 are the observed and expected JC divided by the total number of occupied cells. Therefore, the JCD indicates the difference between the observed and the expected average number of joins (occupied neighbors) per occupied cell, with positive values indicating a higher number of average joins per cell than expected for that occupancy. This allows comparing distributions with differing numbers of presences and in places with differing areas of extent.

#### SAC over time

We examined the temporal change in the SAC of species richness and species distributions by plotting Moran’s I and JCD against the starting year of each atlas period for each region and grain size, respectively. We used linear regressions between JCD and the starting year to assess the overall temporal trend in species distributions for each region and grain size, quantifying shifts in SAC over time independent of occupancy. We also evaluated temporal change in occupancy using linear regressions between the number of occupied cells and the starting year.

#### SAC across spatial grains

To assess the effect of grain size on the SAC of species richness and distributions, we visualized Moran’s I and JCD across different grain sizes for each atlas region and period, utilizing both line plots and box plots.

#### Relationship between change in occupancy and change in SAC

To assess how changes in SAC relate to changes in occupancy, we calculated the log ratios of changes in both metrics by comparing each species’ most recent and earliest records. We modeled the log ratio of the observed JC as a function of the log ratio of occupancy using linear regression. Additionally, we evaluated whether SAC has become more aggregated or dispersed than expected based on changes in occupancy alone. This was done by comparing the observed JC log ratios with expected values derived from the linear regression of the log ratio of expected JC under a random distribution as a function of the log ratio of occupancy. All log ratios were calculated as follows:

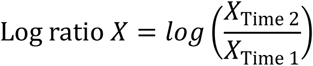

Where 𝑋 represents either occupancy or the observed or expected JC. These log ratios reflect the direction and magnitude of change, with positive values indicating increases and negative values indicating decreases.

All analyses were conducted in R v4.2.3 (R Core Team, 2023) using the following packages: “tidyverse” (v2.0.0; Wickham *et al*., 2019) for data manipulation, “sf” (v1.0-15; Pebesma, 2018; Pebesma & Bivand, 2023) for spatial operations, “spdep” (v1.3-1; Bivand *et al*., 2013) for calculating spatial autocorrelation using the functions “moran.test” and “joincount.test”, and “ggplot2” (v3.4.4; Wickham, 2016) along with “gridExtra” (v2.3; Auguie, 2017) and “viridis” (v0.6.5; Garnier *et al*., 2023) for data visualization.

## Results

### How autocorrelated are species richness and distributions?

#### Species richness

Richness was positively autocorrelated (Moran’s I, p<0.05) across all regions, periods, and grain sizes (Figs. 4a and 5a; Table S1, Supporting Information), with a mean Moran’s I of 0.336 (SD = 0.193). The only exception was Czechia’s coarsest grain size (44x44 km) across all periods, which showed no significant autocorrelation.

**Figure 4.**
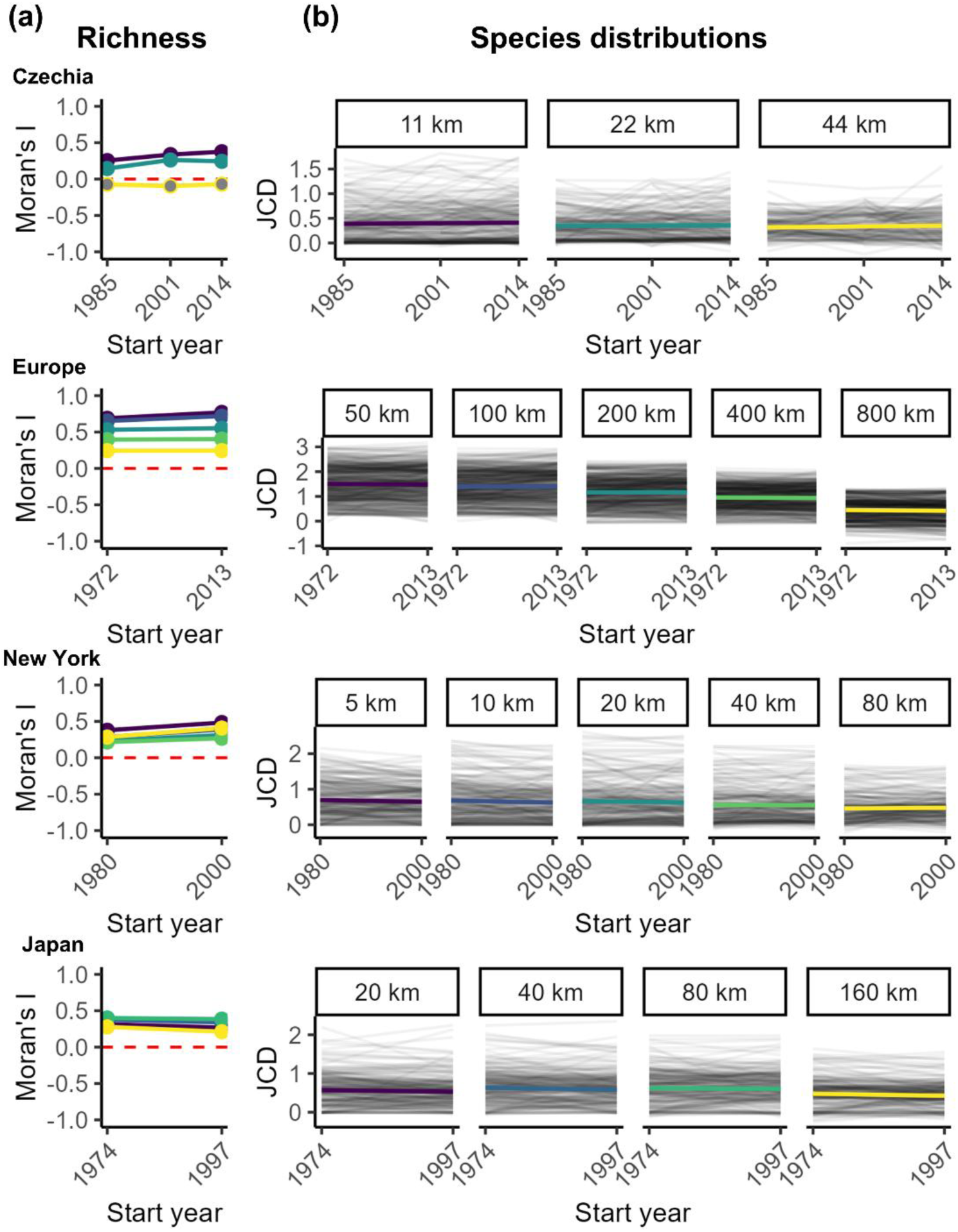
Empirical temporal change in SAC of species diversity and distributions. Panel (a) shows Moran’s I values for richness across atlas periods, with lines connecting values for each grain size. Line colors correspond to grain size and correspond to those in panel (b). Points with a significant (p < 0.05) Moran’s I value are filled with the same color as their respective line, while non-significant points are filled in grey. Panel (b) shows temporal trends in species distributions’ average Join count difference (JCD) across grain sizes (represented by the length of the side of the cell). Trends of individual species are shown in light black lines, while the overall trend for each grain size is shown in color with smoothed regression lines.

**Figure 5.**
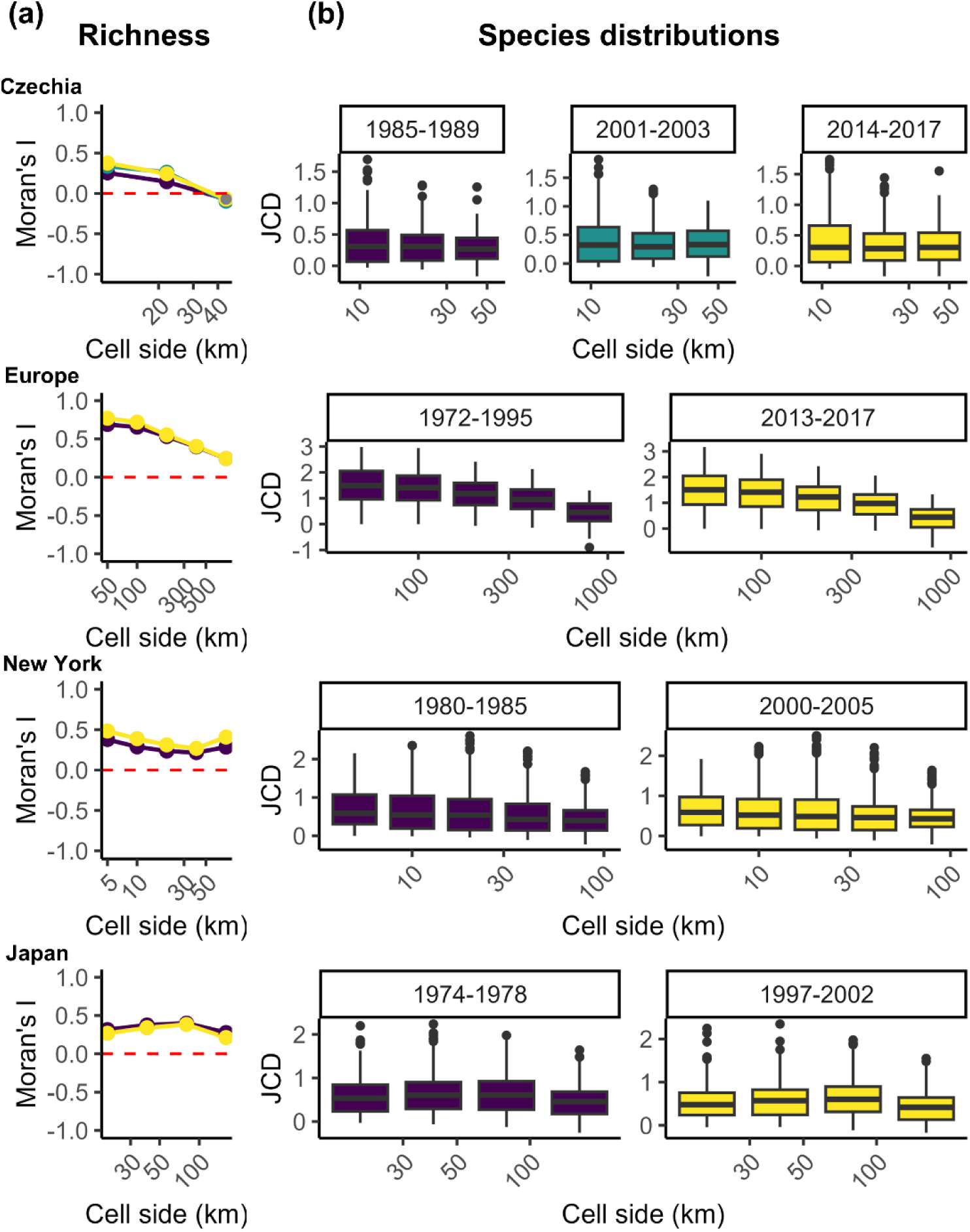
Empirical change in SAC of species diversity and distributions with increasing grain size. Panel (a) illustrates the change in SAC for species richness with increasing grain size (represented by the length of the side of the cell). Moran’s I values are plotted for each grain size and region, with lines and colors representing atlas periods: purple for the first period, yellow for the last, and green for the intermediate when available. Points with significant Moran’s I values (p < 0.05) have the same color fill as their respective line; non-significant points are filled in grey. Panel (b) shows box plots of JCD values for species distributions grouped by grain sizes and atlas periods.

### Species distributions

Most species for which we could calculate JC exhibited significant positive SAC (p < 0.05), ranging from 69.93% in Czechia’s third BBA at 44x44 km to 100% in Europe’s second BBA at 100x100 km (Table S2, Supporting Information), with an overall mean of 88.14% across regions, periods, and grain sizes. JCD values were predominantly positive (Figs. 4b and 5b), indicating most species distributions are positively spatially autocorrelated, independently of their occupancy, across all periods and grain sizes. The percentage of species for which we could calculate the JC ranged from 67.11% in Czechia’s third BBA at 44x44 km and 100% in Europe’s first BBA at 50x50 km (Table S2, Supporting Information).

### SAC over time

#### Species richnessh

Moran’s I of richness remained largely stable over time across all regions and grain sizes, experiencing only minor changes (Fig. 4a; Table S1, Supporting Information). While in Europe and New York, Moran’s I increased over time across all grain sizes, it decreased in Japan and varied in Czechia, showing increases and decreases.

### Species distributions

Individual species distributions exhibited temporal increases, decreases, or no changes in SAC (measured as JCD), across all areas and grain sizes (Fig. 4b). The slopes of JCD over time were generally close to zero (Figs. 4b and 4c) and most were statistically non-significant (Table S3, Supporting Information). Similarly, there was no significant change in occupancy over time (Table S7, Supporting Information).

### SAC across grains

#### Species richness

Moran’s I for species richness declined with increasing grain size in Czechia and Europe (Fig. 5a; Table S1, Supporting Information). In New York, it initially decreased but increased again at the coarsest grain size (80x80 km), while in Japan, Moran’s I first increased and then decreased at the coarsest grain size (160x160 km) in the first two BBAs (Fig. 5a).

### Species distributions

The relationship between JCD and grain size varied across atlas regions (Figs. 5b, 5c; Table S4, Supporting Information). JCD values decreased by more than two-thirds with increasing grain size in Europe. Elsewhere, JCD experienced slight decreases (i.e., New York) or a mix of slight decreases and increases (i.e., Czechia and Japan). The JCD values tended to be higher in Europe than in other regions (between 0.95 and 1.5), and lower in Czechia (between 0.27 and 0.33), while New York and Japan had similar values, between 0.40 and 0.61. All regions showed similar median JCD values at the same grain size across atlas periods (Fig. 5c).

### Relationship between change in occupancy and change in SAC

Temporal changes in SAC were strongly linked to changes in occupancy across regions and grain sizes (all R-squared > 0.79), with most species exhibiting congruent changes in both occupancy and JC across all grain sizes (Figs. 6 and 7; Table S5, Supporting Information). Among species increasing in both occupancy and JC, most had a lower increase in JC than expected (Q4). Similarly, species that declined in both typically showed a lower than expected decrease in JC (Q1). Japan was the exception, with balanced overaggregated (Q1) and overdispersed (Q6) declines in both occupancy and JC, particularly at smaller grain sizes.

**Figure 6.**
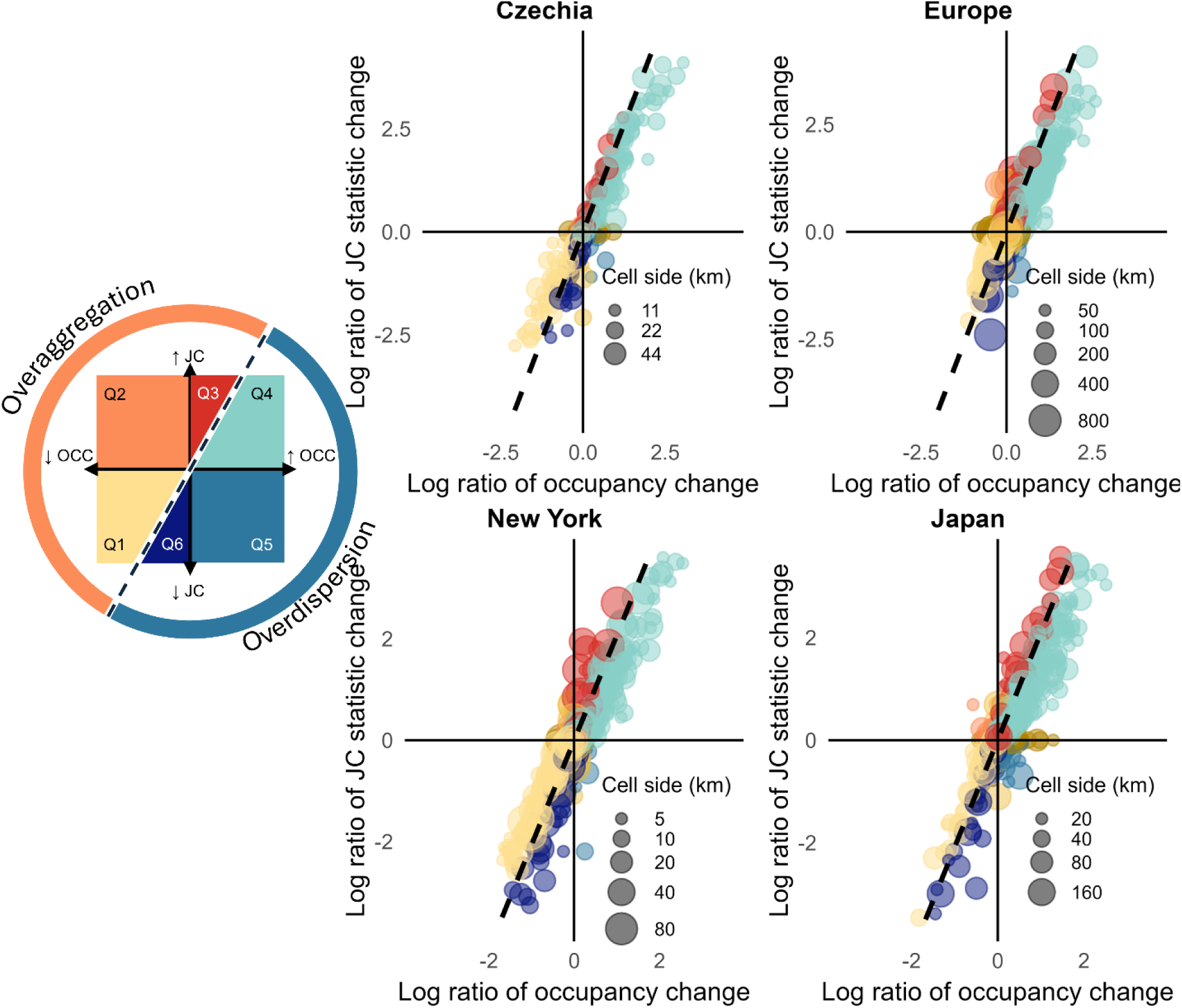
Relationship between empirical occupancy change and Join count statistic (JC) change. The figure shows the log ratio of changes in the observed Join count statistic (JC; y-axis) against the log ratio of changes in occupancy (x-axis) across study areas. Point size corresponds to grain size, with bigger points indicating coarser grains. Colors represent quadrants, categorizing the relationship between JC and occupancy changes, following the conceptual diagram on the left. The black dashed line is the linear regression of the log ratio of expected JC change as a function of occupancy change (i.e., expected JC change for a given occupancy change). Observations to the left of this line indicate a greater-than-expected increase in JC (Q2 and Q3) or a lower-than-expected decrease in JC (Q1) for the observed occupancy change, while observations to the right indicate a lower-than-expected increase in JC (Q4) or a greater-than-expected decrease in JC (Q5 and Q6). Brown observations along the x and y axes are changes in the log ratio of JC independent of changes in occupancy (y-axis) or the reverse (x-axis). No changes in both occupancy and JC and transitions between zero JC and positive JC or the opposite have been removed from the plot for ease of visualization.

**Figure 7.**
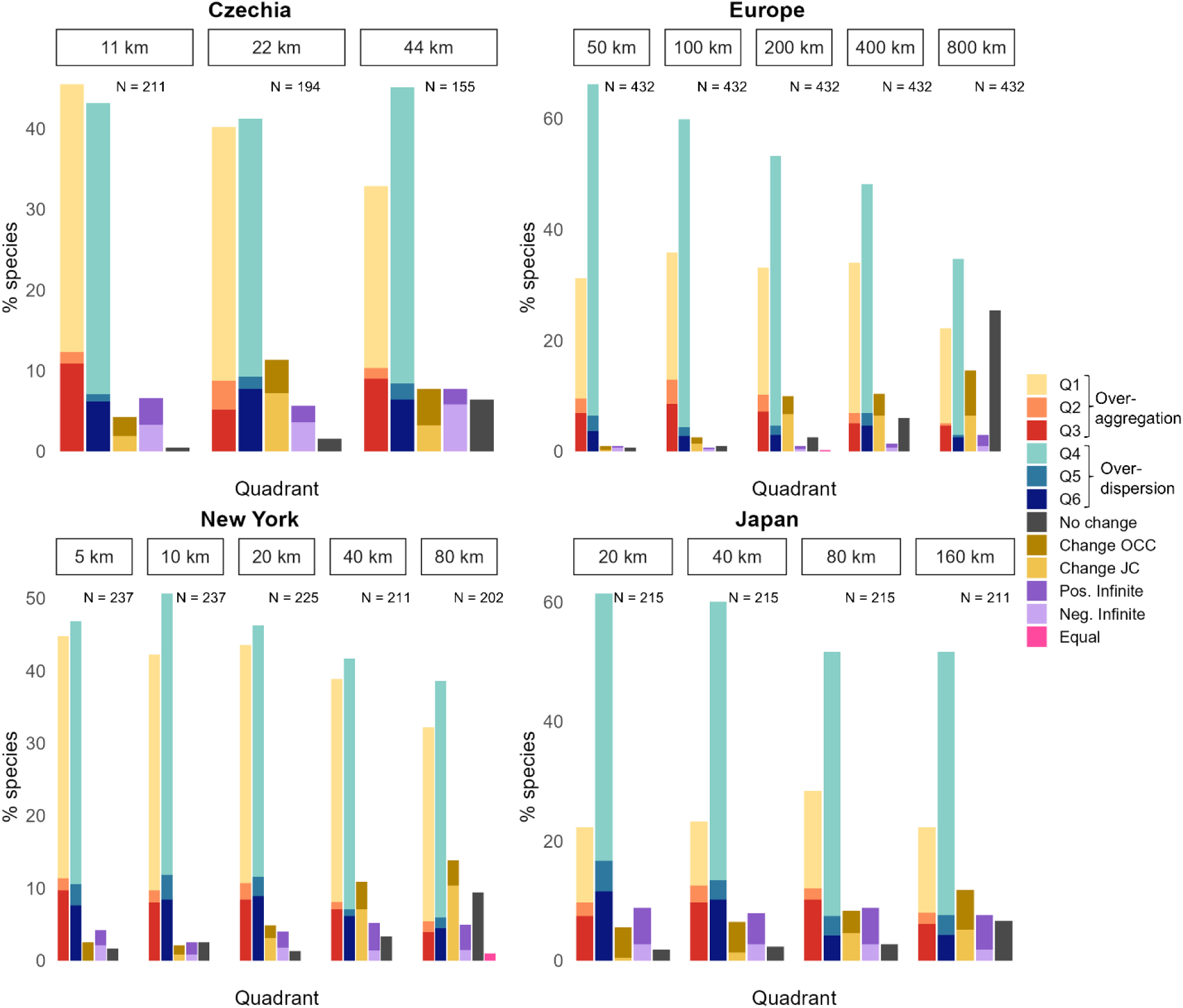
Percentage of species in each category of combined empirical occupancy and Join count statistic (JC) change. Each category (quadrant, Q) of change in the log ratio of the observed occupancy and JC is represented by the same color scheme as the one in Figs. 1b and 6. Quadrants 1, 2, and 3 (yellow, orange, and red) represent cases of overaggregation, where JC decreased less (Q1) or increased more (Q2 and Q3) than expected for the observed change in occupancy. Quadrants 4, 5, and 6 (light blue, mid blue, and dark blue) represent cases of overdispersion, where JC increased less (Q4) or decreased more (Q5 and 6) than expected for the observed change in occupancy. Change OCC represents cases when the distribution experienced a change in occupancy independent of a change in JC, while Change SAC represents changes in JC independent of changes in occupancy. Observations in purple represent transitions between zero JC to positive JC (positive infinite, dark purple) or the reverse (negative infinite, light purple), while those in pink represent observed JC values equal to the expected ones. The total number of species (N) for which we were able to calculate the JC is specified for each area of study and grain size.

The percentage of species in each category of change in occupancy and JC varied with grain size (Fig. 7; Table S6 and Fig. S1, Supporting Information). As spatial grain increased, more species showed no change in both JC and occupancy (No change, Fig. 7), or one of either (Change OCC and Change JC, Fig.7). In contrast, the proportion of species showing both changes in occupancy and JC largely remained stable or decreased (Fig. 7, Fig. S1, Supporting Information).

## Discussion

### Change in SAC and occupancy

As expected, we found a strong positive relationship between changes in occupancy and SAC in species distributions, with increases in occupancy leading to higher SAC and decreases resulting in lower SAC. However, perhaps our most striking finding was that the observed changes in SAC systematically deviated from what we would expect based on changes in occupancy alone. Both expanding and declining species showed less change in SAC than expected: expanding species had lower-than-expected SAC gains, while declining species had lower-than-expected SAC losses.

This suggests that ‘winner’ species (i.e., those with increasing occupancy) are expanding in a disjoint manner and colonizing non-adjacent cells, which contribute fewer spatial joins than more connected cells. We hypothesize this is primarily driven by the high dispersal ability of birds, allowing them to fly over non-suitable habitat patches (Martin & Fahrig, 2018). This pattern remains consistent across scales, likely reflecting a heavy-tailed dispersal distribution, where most individuals move short distances while a few disperse over long distances, a phenomenon common in European breeding birds (Fandos et al., 2023).

Conversely, ‘loser’ species (i.e., those with decreasing occupancy) are being extirpated in isolated or less connected cells, leaving the remaining cells more spatially clustered. This aligns with the ‘rescue effect’, where extirpations are more frequent in disjoint areas that receive fewer dispersing individuals (Hanski, 1999). Howard et al. (2023) also observed this pattern in the European BBAs, finding that extirpations were more likely in distant cells, while colonizations occurred closer to continually occupied grid cells. We attribute the discrepancy in colonization patterns to the fact that we are focusing on immediately adjacent cells, while Howard et al. (2023) considered the distance between cells. Thus, isolated populations appear to be key drivers of range dynamics, particularly dynamics of spatial aggregation of species distributions. So far, we are unaware of these patterns being empirically described in the literature.

Despite observing variation in changes in the JCD across individual species, we found zero mean temporal trends in JCD or occupancy when averaging across species. This zero mean, but large variation, is similar to the results of previous studies that found zero average net temporal change in local-scale metrics like richness (Vellend et al. 2013; Dornelas et al. 2014, 2023; Blowes et al. 2019; Crockett et al. 2022), despite there being many increases, decreases, and changes in turnover (Dornelas et al. 2014, 2023; Blowes et al. 2019). Another example of this phenomenon is the relatively weak average global decline in population sizes, coupled with some extreme losses (Leung et al. 2020). All this indicates that we should perhaps shift focus from average trends to the variation around the mean. In our case, despite the overall zero net trend in SAC, some species still showed increases and decreases in both their SAC and occupancy, providing a starting point for identifying species for which we should prioritize conservation actions in follow-up studies.

Furthermore, while we observed zero average changes in occupancy and SAC independently, their joint relationship revealed deviations from the expected dynamics, providing deeper insights into species biodiversity change and potential conservation needs. For instance, although an increase in autocorrelation in the range of a species suggests a positive effect of a decrease in fragmentation, this could be due to a range contraction to a single or few patches. This emphasizes the importance of jointly considering multiple metrics when studying biodiversity change, something already pointed out by other studies. For instance, Blowes et al. (2024) combined alpha and gamma diversity to study community homogenization and differentiation; Hillebrand et al. (2018) advocated for considering multiple aspects, such as changes in species identity and dominance, rather than uniquely focusing on species richness; and Blowes et al. (2022) recommended using a combination of abundance, evenness, and richness. Our findings further reinforce this idea.

#### SAC, grain, and extent

Most species distributions exhibited significant positive SAC across all regions, time periods, and grain sizes, indicating that spatial clustering is indeed a ubiquitous feature of distributions (Legendre 1993). However, we observed regional differences in how median species SAC changed with grain size.

Median JCD showed very slight changes across grain sizes in Czechia, New York, and Japan. This potentially indicates that the spatial processes driving distributions are consistent across these spatial scales (between 5x5 and 160x160 km). Moreover, these regions had similar median JCD values and trends across grains. This supports the robustness of our findings, particularly as they share observations with comparable grain sizes.

Europe, with the largest spatial extent, exhibited the highest median SAC values, even at grain sizes similar to those of other areas. Here, median SAC decreased with increasing grain size, consistent with Hui’s (2009) findings. This trend likely reflects the loss of finer-scale spatial structure as species presences become aggregated over larger sampling units. Larger grains typically encompass a wider range of habitats, which can dilute SAC by averaging across diverse environmental conditions and reducing spatial differentiation. Although less pronounced, a similar decreasing trend in JCD with grain size was also observed in the other regions.

Europe’s higher SAC values may be attributed to its larger extent, which captures the entire distributional range of many species and a broader environmental variability. This is not simply due to cell number—New York consistently had the most cells at each grain size (Table S8, Supporting Information), yet had lower SAC values. In addition to extent, the position of the atlas within the full range of a species may also influence the observed patterns of range expansion or contraction. We suggest that future work should explore how geographic extent affects SAC metrics, and perhaps better null models can be designed.

The observed scale dependency of SAC has important implications for species distribution modeling (SDMs) and broader ecological research. Stronger SAC at finer grains suggests species distributions may be more predictable at these scales, enabling more accurate spatial and temporal interpolations (Keil & Chase 2019, 2022). However, SAC may also introduce biases in SDMs if not accounted for, or may signal that a relevant driver of distributions is being overlooked. Coarser resolutions can reduce these biases, but at the cost of losing information (Boyd *et al*., 2024). These findings underscore the need to consider grain size and spatial extent carefully when interpreting SAC and its underlying drivers in ecological studies.

#### Species diversity

Trends in the SAC of richness across time and grain sizes mirrored the patterns observed in species distributions. This is unsurprising, as species richness represents the aggregate of individual species distributions (Gaston, 2000; Hawkins, 2012). Thus, changes in the spatial structure of species distributions will be reflected in the spatial structure of richness. The similar patterns revealed by both Moran’s I and JC metrics highlight the robustness of our results, confirming that the observed spatial autocorrelation is independent of the metric used.

#### Challenges and pitfalls

We acknowledge several limitations to our work. First, unknown sampling effort and imperfect detection could influence the observed patterns, particularly at finer scales, where variations in these metrics can have a greater impact (Boyd *et al*., 2024). While methods such as the Frescalo algorithm (Hill, 2012) or occupancy models account for this by incorporating spatial autocorrelation or modeling it as part of the observation process or the species’ distributions (MacKenzie *et al*., 2018), this approach is challenging for studies like ours, which specifically focus on spatial autocorrelation itself. In other words, we cannot use the existing toolbox to account for sampling effort to study autocorrelation because these methods introduce additional autocorrelation. Nonetheless, the issue of unknown sampling effort and detectability should become less pronounced at coarser scales (Hurlbert & Jetz, 2007; Keil *et al*., 2014), making our results more robust as grain size increases. Furthermore, if the sampling process and detection are random at fine grains, they are more likely to reduce observed SAC at fine grains rather than inflate it. Thus, the observed patterns, particularly the increase of SAC towards fine grain size, are likely genuine.

Second, global SAC metrics, such as the ones we used, may obscure finer-scale, localized patterns of SAC and the ecological and environmental processes that vary across space (Rollinson *et al*., 2021). Very different distribution configurations can have the same global SAC value. Therefore, spatially explicit maps of local SAC can provide a more informative summary of species distributions, particularly when monitored in time. Third, here we focused on adjacency-based SAC, while distance-based SAC can provide better insights into population connectivity. Fourth, we did not consider invasive species separately, although they might be experiencing different spatial autocorrelation patterns, and, indeed, SAC has been used to study invasion processes (Wang et al., 2011). Lastly, there is a geographical bias, as our study is focused on areas in the Northern Hemisphere, however, the observed patterns can be different in other areas.

#### Future pathways

We recommend further exploration of SAC as a biodiversity metric and support using multiple metrics to better understand and characterize biodiversity temporal dynamics. Follow-up studies should investigate local SAC dynamics using local metrics such as LISA (Anselin, 1995) or LICD (Anselin & Li, 2019) to identify and track high and low SAC hotspots over time. Incorporating scale-dependent drivers—such as climate and land use—and examining the scales at which these drivers exhibit spatial autocorrelation and their alignment with distribution SAC patterns could provide deeper insights into the ecological and environmental processes shaping biodiversity across scales.

Additionally, exploring the SAC of sampling effort and comparing these results with ours could highlight its influence on observed patterns. Extending analyses to datasets with different taxa, geographic areas, and grain sizes would help assess the generality of our findings. Finally, investigating distance-based SAC and identifying the distance thresholds at which species distributions remain autocorrelated could yield valuable insights into connectivity, as well as advance the identification of the spatial extent to which biodiversity data can be interpolated in under-sampled regions.

## Data availability statement

All raw atlas data are available by request from the respective atlas coordinators. Codes, derived data used in the analyses, and results are available in the Zenodo repository (https://doi.org/10.5281/zenodo.15463598).

