## supporting information for "Spatial autocorrelation of species diversity and distributions in time and across spatial scales"

**Breeding Bird Atlases (BBAs) data preparation**

We only considered full species and removed hybrids and subspecies from our data. We also removed cells that had no landmass (fully aquatic cells).

The Japanese BBAs followed two sampling strategies: questionnaires and fieldwork. We were not able to distinguish the sampling strategy for each data point in the first two BBAs. Therefore, even though we could filter by strategy in the second BBA, we chose to keep both, ensuring both BBAs were comparable. The third Japanese BBA had a substantially higher survey effort, evidenced by the addition of 147 species that had not been observed in the previous two atlases. We therefore chose to only use the first two BBAs.

In the European BBAs, we only considered observations classified as “Apparent trust” and “Trusted occurrence”. The first European BBA was conducted between 1972 and 1995, however, most of its data was collected during the 1980s. To compare both European BBA periods, we restricted the area of study to a well-defined geographical area based on the coverage of the first BBA (Keller et al., 2020). This excluded Cyprus, the Asian part of Turkey, and the Canary Islands (not included in the first BBA), as well as most of European Russia, Kazakhstan, Georgia, Armenia, and Azerbaijan (partially surveyed in the first BBA). Cells that lied totally or mostly (>70% of their area) within the above-mentioned areas were not included. Additionally, 70 grid cells were not assigned exactly to the same land areas in the first and second BBA, as the first BBA merged neighboring grids while the second did not. We used a version of the second BBA grid that had these cells merged.

**Table S1. How autocorrelated is richness?**

| **Study area** | **Period** | **Cell side** | **Moran’s I** | **p-value** |
| --- | --- | --- | --- | --- |
| Czechia | 1985-1989 | 11 | 0.254 | <0.01 |
| Czechia | 1985-1989 | 22 | 0.145 | <0.01 |
| Czechia | 1985-1989 | 44 | -0.073 | 0.77 |
| Czechia | 2001-2003 | 11 | 0.336 | <0.01 |
| Czechia | 2001-2003 | 22 | 0.263 | <0.01 |
| Czechia | 2001-2003 | 44 | -0.096 | 0.837 |
| Czechia | 2014-2017 | 11 | 0.376 | <0.01 |
| Czechia | 2014-2017 | 22 | 0.244 | <0.01 |
| Czechia | 2014-2017 | 44 | -0.07 | 0.742 |
| Europe | 1972-1995 | 50 | 0.689 | <0.01 |
| Europe | 1972-1995 | 100 | 0.653 | <0.01 |
| Europe | 1972-1995 | 200 | 0.531 | <0.01 |
| Europe | 1972-1995 | 400 | 0.396 | <0.01 |
| Europe | 1972-1995 | 800 | 0.244 | 0.021 |
| Europe | 2013-2017 | 50 | 0.77 | <0.01 |
| Europe | 2013-2017 | 100 | 0.719 | <0.01 |
| Europe | 2013-2017 | 200 | 0.553 | <0.01 |
| Europe | 2013-2017 | 400 | 0.404 | <0.01 |
| Europe | 2013-2017 | 800 | 0.245 | 0.02 |
| New York | 1980-1985 | 5 | 0.378 | <0.01 |
| New York | 1980-1985 | 10 | 0.288 | <0.01 |
| New York | 1980-1985 | 20 | 0.235 | <0.01 |
| New York | 1980-1985 | 40 | 0.216 | <0.01 |
| New York | 1980-1985 | 80 | 0.284 | <0.01 |
| New York | 2000-2005 | 5 | 0.484 | <0.01 |
| New York | 2000-2005 | 10 | 0.389 | <0.01 |
| New York | 2000-2005 | 20 | 0.312 | <0.01 |
| New York | 2000-2005 | 40 | 0.269 | <0.01 |
| New York | 2000-2005 | 80 | 0.409 | <0.01 |
| Japan | 1974-1978 | 20 | 0.316 | <0.01 |
| Japan | 1974-1978 | 40 | 0.377 | <0.01 |
| Japan | 1974-1978 | 80 | 0.401 | <0.01 |
| Japan | 1974-1978 | 160 | 0.278 | <0.01 |
| Japan | 1997-2002 | 20 | 0.269 | <0.01 |
| Japan | 1997-2002 | 40 | 0.34 | <0.01 |
| Japan | 1997-2002 | 80 | 0.384 | <0.01 |
| Japan | 1997-2002 | 160 | 0.212 | 0.019 |

**Table S2. How autocorrelated are species distributions?**

| **Study area** | **Period** | **Cell side** | **N species** | **N species SAC** | **N species positive SAC** | **% species positive SAC** |
| --- | --- | --- | --- | --- | --- | --- |
| Czechia | 1985-1989 | 11 | 206 | 201 | 181 | 90.05 |
| Czechia | 1985-1989 | 22 | 206 | 184 | 173 | 94.02 |
| Czechia | 1985-1989 | 44 | 206 | 173 | 122 | 70.52 |
| Czechia | 2001-2003 | 11 | 213 | 206 | 173 | 83.98 |
| Czechia | 2001-2003 | 22 | 213 | 187 | 171 | 91.44 |
| Czechia | 2001-2003 | 44 | 213 | 145 | 103 | 71.03 |
| Czechia | 2014-2017 | 11 | 228 | 215 | 191 | 88.84 |
| Czechia | 2014-2017 | 22 | 228 | 197 | 173 | 87.82 |
| Czechia | 2014-2017 | 44 | 228 | 153 | 107 | 69.93 |
| Europe | 1972-1995 | 50 | 432 | 432 | 431 | 99.77 |
| Europe | 1972-1995 | 100 | 432 | 431 | 431 | 100 |
| Europe | 1972-1995 | 200 | 432 | 430 | 424 | 98.6 |
| Europe | 1972-1995 | 400 | 432 | 428 | 415 | 96.96 |
| Europe | 1972-1995 | 800 | 432 | 384 | 279 | 72.66 |
| Europe | 2013-2017 | 50 | 446 | 444 | 440 | 99.1 |
| Europe | 2013-2017 | 100 | 446 | 444 | 441 | 99.32 |
| Europe | 2013-2017 | 200 | 446 | 443 | 433 | 97.74 |
| Europe | 2013-2017 | 400 | 446 | 439 | 421 | 95.9 |
| Europe | 2013-2017 | 800 | 446 | 381 | 273 | 71.65 |
| New York | 1980-1985 | 5 | 242 | 236 | 225 | 95.34 |
| New York | 1980-1985 | 10 | 242 | 236 | 226 | 95.76 |
| New York | 1980-1985 | 20 | 242 | 227 | 210 | 92.51 |
| New York | 1980-1985 | 40 | 242 | 211 | 185 | 87.68 |
| New York | 1980-1985 | 80 | 242 | 205 | 149 | 72.68 |
| New York | 2000-2005 | 5 | 248 | 242 | 232 | 95.87 |
| New York | 2000-2005 | 10 | 248 | 240 | 231 | 96.25 |
| New York | 2000-2005 | 20 | 248 | 232 | 218 | 93.97 |
| New York | 2000-2005 | 40 | 248 | 222 | 198 | 89.19 |
| New York | 2000-2005 | 80 | 248 | 207 | 165 | 79.71 |
| Japan | 1974-1978 | 20 | 226 | 218 | 193 | 88.53 |
| Japan | 1974-1978 | 40 | 226 | 218 | 193 | 88.53 |
| Japan | 1974-1978 | 80 | 226 | 214 | 178 | 83.18 |
| Japan | 1974-1978 | 160 | 226 | 205 | 161 | 78.54 |
| Japan | 1997-2002 | 20 | 239 | 229 | 207 | 90.39 |
| Japan | 1997-2002 | 40 | 241 | 227 | 202 | 88.99 |
| Japan | 1997-2002 | 80 | 241 | 223 | 197 | 88.34 |
| Japan | 1997-2002 | 160 | 243 | 217 | 166 | 76.5 |

**Table S3. SAC change over time**

| **Study area** | **CellSide** | **JCD slope** | **SE** | **p-value** | **R-squared** |
| --- | --- | --- | --- | --- | --- |
| Czechia | 11 | 0.0008 | 0.0013 | 0.543 | 0.0006 |
| Czechia | 22 | 0.0002 | 0.0011 | 0.851 | 0.0001 |
| Czechia | 44 | 0.00117 | 0.001 | 0.237 | 0.0029 |
| Europe | 50 | -0.00073 | 0.0011 | 0.526 | 0.0005 |
| Europe | 100 | 0.00004 | 0.0011 | 0.967 | >0.0001 |
| Europe | 200 | 0.0002 | 0.001 | 0.840 | >0.0001 |
| Europe | 400 | -0.00043 | 0.0008 | 0.594 | 0.0003 |
| Europe | 800 | -0.00078 | 0.0007 | 0.284 | 0.0013 |
| New York | 5 | -0.00237 | 0.0022 | 0.279 | 0.0023 |
| New York | 10 | -0.0028 | 0.0025 | 0.261 | 0.0026 |
| New York | 20 | -0.00203 | 0.0028 | 0.464 | 0.0011 |
| New York | 40 | -0.00064 | 0.0024 | 0.795 | 0.0002 |
| New York | 80 | 0.00096 | 0.0019 | 0.611 | 0.0006 |
| Japan | 20 | -0.00184 | 0.0017 | 0.280 | 0.0025 |
| Japan | 40 | -0.00222 | 0.0018 | 0.219 | 0.0033 |
| Japan | 80 | -0.00081 | 0.0018 | 0.646 | 0.0005 |
| Japan | 160 | -0.00224 | 0.0015 | 0.133 | 0.0049 |

**Table S4. Median Join count difference across grain sizes**

| **Study area** | **Period** | **Grain size** | **Median JCD** |
| --- | --- | --- | --- |
| Czechia | 1985-1989 | 11 | 0.308283 |
| Czechia | 1985-1989 | 22 | 0.307145 |
| Czechia | 1985-1989 | 44 | 0.268037 |
| Czechia | 2001-2003 | 11 | 0.322621 |
| Czechia | 2001-2003 | 22 | 0.293658 |
| Czechia | 2001-2003 | 44 | 0.331062 |
| Czechia | 2014-2017 | 11 | 0.303721 |
| Czechia | 2014-2017 | 22 | 0.283699 |
| Czechia | 2014-2017 | 44 | 0.303484 |
| Europe | 1972-1995 | 50 | 1.48875 |
| Europe | 1972-1995 | 100 | 1.40859 |
| Europe | 1972-1995 | 200 | 1.175764 |
| Europe | 1972-1995 | 400 | 0.952956 |
| Europe | 1972-1995 | 800 | 0.454414 |
| Europe | 2013-2017 | 50 | 1.508319 |
| Europe | 2013-2017 | 100 | 1.415767 |
| Europe | 2013-2017 | 200 | 1.22958 |
| Europe | 2013-2017 | 400 | 0.977564 |
| Europe | 2013-2017 | 800 | 0.446408 |
| New York | 1980-1985 | 5 | 0.586071 |
| New York | 1980-1985 | 10 | 0.543371 |
| New York | 1980-1985 | 20 | 0.537822 |
| New York | 1980-1985 | 40 | 0.428573 |
| New York | 1980-1985 | 80 | 0.398649 |
| New York | 2000-2005 | 5 | 0.589625 |
| New York | 2000-2005 | 10 | 0.520527 |
| New York | 2000-2005 | 20 | 0.486873 |
| New York | 2000-2005 | 40 | 0.459942 |
| New York | 2000-2005 | 80 | 0.43018 |
| Japan | 1974-1978 | 20 | 0.534285 |
| Japan | 1974-1978 | 40 | 0.604162 |
| Japan | 1974-1978 | 80 | 0.608167 |
| Japan | 1974-1978 | 160 | 0.456815 |
| Japan | 1997-2002 | 20 | 0.478103 |
| Japan | 1997-2002 | 40 | 0.570985 |
| Japan | 1997-2002 | 80 | 0.601907 |
| Japan | 1997-2002 | 160 | 0.41626 |

**Table S5. Summary of linear models of log ratio of observed change in join count and occupancy**

| **Study area** | **Grain size** | **Slope** | **p-value** | **R-squared** |
| --- | --- | --- | --- | --- |
| Czechia | 11 | 1.467442 | <0.01 | 0.910212 |
| Czechia | 22 | 1.485308 | <0.01 | 0.890957 |
| Czechia | 44 | 1.714557 | <0.01 | 0.91236 |
| New York | 5 | 1.443444 | <0.01 | 0.904832 |
| New York | 10 | 1.534466 | <0.01 | 0.864794 |
| New York | 20 | 1.589218 | <0.01 | 0.894493 |
| New York | 40 | 1.642021 | <0.01 | 0.876472 |
| New York | 80 | 1.786007 | <0.01 | 0.884027 |
| Europe | 50 | 1.305308 | <0.01 | 0.903208 |
| Europe | 100 | 1.394025 | <0.01 | 0.916779 |
| Europe | 200 | 1.511387 | <0.01 | 0.890615 |
| Europe | 400 | 1.545626 | <0.01 | 0.862787 |
| Europe | 800 | 1.571874 | <0.01 | 0.791872 |
| Japan | 20 | 1.419799 | <0.01 | 0.814283 |
| Japan | 40 | 1.537705 | <0.01 | 0.874545 |
| Japan | 80 | 1.634504 | <0.01 | 0.847362 |
| Japan | 160 | 1.693378 | <0.01 | 0.843724 |

**Table S6. Percentage of species in each quadrant of the combined change in occupancy and JC per atlas location and grain size**

| **Study area** | **Grain size** | **Quadrant** | **N species** | **Total species log ratio** | **% species** |
| --- | --- | --- | --- | --- | --- |
| Czechia | 11 | Change OCC | 5 | 211 | 2.369668 |
| Czechia | 11 | Change JC | 4 | 211 | 1.895735 |
| Czechia | 11 | Neg. Infinite | 7 | 211 | 3.317536 |
| Czechia | 11 | No change | 1 | 211 | 0.473934 |
| Czechia | 11 | Pos. Infinite | 7 | 211 | 3.317536 |
| Czechia | 11 | Q1 | 70 | 211 | 33.17536 |
| Czechia | 11 | Q2 | 3 | 211 | 1.421801 |
| Czechia | 11 | Q3 | 23 | 211 | 10.90047 |
| Czechia | 11 | Q4 | 76 | 211 | 36.01896 |
| Czechia | 11 | Q5 | 2 | 211 | 0.947867 |
| Czechia | 11 | Q6 | 13 | 211 | 6.161137 |
| Czechia | 22 | Change OCC | 8 | 194 | 4.123711 |
| Czechia | 22 | Change JC | 14 | 194 | 7.216495 |
| Czechia | 22 | Neg. Infinite | 7 | 194 | 3.608247 |
| Czechia | 22 | No change | 3 | 194 | 1.546392 |
| Czechia | 22 | Pos. Infinite | 4 | 194 | 2.061856 |
| Czechia | 22 | Q1 | 61 | 194 | 31.4433 |
| Czechia | 22 | Q2 | 7 | 194 | 3.608247 |
| Czechia | 22 | Q3 | 10 | 194 | 5.154639 |
| Czechia | 22 | Q4 | 62 | 194 | 31.95876 |
| Czechia | 22 | Q5 | 3 | 194 | 1.546392 |
| Czechia | 22 | Q6 | 15 | 194 | 7.731959 |
| Czechia | 44 | Change OCC | 7 | 155 | 4.516129 |
| Czechia | 44 | Change JC | 5 | 155 | 3.225806 |
| Czechia | 44 | Neg. Infinite | 9 | 155 | 5.806452 |
| Czechia | 44 | No change | 10 | 155 | 6.451613 |
| Czechia | 44 | Pos. Infinite | 3 | 155 | 1.935484 |
| Czechia | 44 | Q1 | 35 | 155 | 22.58065 |
| Czechia | 44 | Q2 | 2 | 155 | 1.290323 |
| Czechia | 44 | Q3 | 14 | 155 | 9.032258 |
| Czechia | 44 | Q4 | 57 | 155 | 36.77419 |
| Czechia | 44 | Q5 | 3 | 155 | 1.935484 |
| Czechia | 44 | Q6 | 10 | 155 | 6.451613 |
| Europe | 50 | Change OCC | 3 | 432 | 0.694444 |
| Europe | 50 | Change JC | 1 | 432 | 0.231481 |
| Europe | 50 | Neg. Infinite | 3 | 432 | 0.694444 |
| Europe | 50 | No change | 3 | 432 | 0.694444 |
| Europe | 50 | Pos. Infinite | 1 | 432 | 0.231481 |
| Europe | 50 | Q1 | 94 | 432 | 21.75926 |
| Europe | 50 | Q2 | 11 | 432 | 2.546296 |
| Europe | 50 | Q3 | 30 | 432 | 6.944444 |
| Europe | 50 | Q4 | 258 | 432 | 59.72222 |
| Europe | 50 | Q5 | 12 | 432 | 2.777778 |
| Europe | 50 | Q6 | 16 | 432 | 3.703704 |
| Europe | 100 | Change OCC | 5 | 432 | 1.157407 |
| Europe | 100 | Change JC | 6 | 432 | 1.388889 |
| Europe | 100 | Neg. Infinite | 2 | 432 | 0.462963 |
| Europe | 100 | No change | 4 | 432 | 0.925926 |
| Europe | 100 | Pos. Infinite | 1 | 432 | 0.231481 |
| Europe | 100 | Q1 | 99 | 432 | 22.91667 |
| Europe | 100 | Q2 | 19 | 432 | 4.398148 |
| Europe | 100 | Q3 | 37 | 432 | 8.564815 |
| Europe | 100 | Q4 | 240 | 432 | 55.55556 |
| Europe | 100 | Q5 | 7 | 432 | 1.62037 |
| Europe | 100 | Q6 | 12 | 432 | 2.777778 |
| Europe | 200 | Change OCC | 14 | 432 | 3.240741 |
| Europe | 200 | Change JC | 29 | 432 | 6.712963 |
| Europe | 200 | Equal | 1 | 432 | 0.231481 |
| Europe | 200 | Neg. Infinite | 2 | 432 | 0.462963 |
| Europe | 200 | No change | 11 | 432 | 2.546296 |
| Europe | 200 | Pos. Infinite | 2 | 432 | 0.462963 |
| Europe | 200 | Q1 | 99 | 432 | 22.91667 |
| Europe | 200 | Q2 | 13 | 432 | 3.009259 |
| Europe | 200 | Q3 | 31 | 432 | 7.175926 |
| Europe | 200 | Q4 | 210 | 432 | 48.61111 |
| Europe | 200 | Q5 | 7 | 432 | 1.62037 |
| Europe | 200 | Q6 | 13 | 432 | 3.009259 |
| Europe | 400 | Change OCC | 17 | 432 | 3.935185 |
| Europe | 400 | Change JC | 28 | 432 | 6.481481 |
| Europe | 400 | Neg. Infinite | 3 | 432 | 0.694444 |
| Europe | 400 | No change | 26 | 432 | 6.018519 |
| Europe | 400 | Pos. Infinite | 3 | 432 | 0.694444 |
| Europe | 400 | Q1 | 117 | 432 | 27.08333 |
| Europe | 400 | Q2 | 8 | 432 | 1.851852 |
| Europe | 400 | Q3 | 22 | 432 | 5.092593 |
| Europe | 400 | Q4 | 178 | 432 | 41.2037 |
| Europe | 400 | Q5 | 10 | 432 | 2.314815 |
| Europe | 400 | Q6 | 20 | 432 | 4.62963 |
| Europe | 800 | Change OCC | 35 | 432 | 8.101852 |
| Europe | 800 | Change JC | 28 | 432 | 6.481481 |
| Europe | 800 | Neg. Infinite | 4 | 432 | 0.925926 |
| Europe | 800 | No change | 110 | 432 | 25.46296 |
| Europe | 800 | Pos. Infinite | 9 | 432 | 2.083333 |
| Europe | 800 | Q1 | 74 | 432 | 17.12963 |
| Europe | 800 | Q2 | 2 | 432 | 0.462963 |
| Europe | 800 | Q3 | 20 | 432 | 4.62963 |
| Europe | 800 | Q4 | 137 | 432 | 31.71296 |
| Europe | 800 | Q5 | 2 | 432 | 0.462963 |
| Europe | 800 | Q6 | 11 | 432 | 2.546296 |
| New York | 5 | Change OCC | 6 | 237 | 2.531646 |
| New York | 5 | Neg. Infinite | 5 | 237 | 2.109705 |
| New York | 5 | No change | 4 | 237 | 1.687764 |
| New York | 5 | Pos. Infinite | 5 | 237 | 2.109705 |
| New York | 5 | Q1 | 79 | 237 | 33.33333 |
| New York | 5 | Q2 | 4 | 237 | 1.687764 |
| New York | 5 | Q3 | 23 | 237 | 9.704641 |
| New York | 5 | Q4 | 86 | 237 | 36.28692 |
| New York | 5 | Q5 | 7 | 237 | 2.953586 |
| New York | 5 | Q6 | 18 | 237 | 7.594937 |
| New York | 10 | Change OCC | 3 | 237 | 1.265823 |
| New York | 10 | Change JC | 2 | 237 | 0.843882 |
| New York | 10 | Neg. Infinite | 2 | 237 | 0.843882 |
| New York | 10 | No change | 6 | 237 | 2.531646 |
| New York | 10 | Pos. Infinite | 4 | 237 | 1.687764 |
| New York | 10 | Q1 | 77 | 237 | 32.48945 |
| New York | 10 | Q2 | 4 | 237 | 1.687764 |
| New York | 10 | Q3 | 19 | 237 | 8.016878 |
| New York | 10 | Q4 | 92 | 237 | 38.81857 |
| New York | 10 | Q5 | 8 | 237 | 3.375527 |
| New York | 10 | Q6 | 20 | 237 | 8.438819 |
| New York | 20 | Change OCC | 4 | 225 | 1.777778 |
| New York | 20 | Change JC | 7 | 225 | 3.111111 |
| New York | 20 | Neg. Infinite | 4 | 225 | 1.777778 |
| New York | 20 | No change | 3 | 225 | 1.333333 |
| New York | 20 | Pos. Infinite | 5 | 225 | 2.222222 |
| New York | 20 | Q1 | 74 | 225 | 32.88889 |
| New York | 20 | Q2 | 5 | 225 | 2.222222 |
| New York | 20 | Q3 | 19 | 225 | 8.444444 |
| New York | 20 | Q4 | 78 | 225 | 34.66667 |
| New York | 20 | Q5 | 6 | 225 | 2.666667 |
| New York | 20 | Q6 | 20 | 225 | 8.888889 |
| New York | 40 | Change OCC | 8 | 211 | 3.791469 |
| New York | 40 | Change JC | 15 | 211 | 7.109005 |
| New York | 40 | Neg. Infinite | 3 | 211 | 1.421801 |
| New York | 40 | No change | 7 | 211 | 3.317536 |
| New York | 40 | Pos. Infinite | 8 | 211 | 3.791469 |
| New York | 40 | Q1 | 65 | 211 | 30.80569 |
| New York | 40 | Q2 | 2 | 211 | 0.947867 |
| New York | 40 | Q3 | 15 | 211 | 7.109005 |
| New York | 40 | Q4 | 73 | 211 | 34.59716 |
| New York | 40 | Q5 | 2 | 211 | 0.947867 |
| New York | 40 | Q6 | 13 | 211 | 6.161137 |
| New York | 80 | Change OCC | 7 | 202 | 3.465347 |
| New York | 80 | Change JC | 21 | 202 | 10.39604 |
| New York | 80 | Equal | 2 | 202 | 0.990099 |
| New York | 80 | Neg. Infinite | 3 | 202 | 1.485149 |
| New York | 80 | No change | 19 | 202 | 9.405941 |
| New York | 80 | Pos. Infinite | 7 | 202 | 3.465347 |
| New York | 80 | Q1 | 54 | 202 | 26.73267 |
| New York | 80 | Q2 | 3 | 202 | 1.485149 |
| New York | 80 | Q3 | 8 | 202 | 3.960396 |
| New York | 80 | Q4 | 66 | 202 | 32.67327 |
| New York | 80 | Q5 | 3 | 202 | 1.485149 |
| New York | 80 | Q6 | 9 | 202 | 4.455446 |
| Japan | 20 | Change OCC | 11 | 215 | 5.116279 |
| Japan | 20 | Change JC | 1 | 215 | 0.465116 |
| Japan | 20 | Neg. Infinite | 6 | 215 | 2.790698 |
| Japan | 20 | No change | 4 | 215 | 1.860465 |
| Japan | 20 | Pos. Infinite | 13 | 215 | 6.046512 |
| Japan | 20 | Q1 | 27 | 215 | 12.55814 |
| Japan | 20 | Q2 | 5 | 215 | 2.325581 |
| Japan | 20 | Q3 | 16 | 215 | 7.44186 |
| Japan | 20 | Q4 | 96 | 215 | 44.65116 |
| Japan | 20 | Q5 | 11 | 215 | 5.116279 |
| Japan | 20 | Q6 | 25 | 215 | 11.62791 |
| Japan | 40 | Change OCC | 11 | 215 | 5.116279 |
| Japan | 40 | Change JC | 3 | 215 | 1.395349 |
| Japan | 40 | Neg. Infinite | 6 | 215 | 2.790698 |
| Japan | 40 | No change | 5 | 215 | 2.325581 |
| Japan | 40 | Pos. Infinite | 11 | 215 | 5.116279 |
| Japan | 40 | Q1 | 23 | 215 | 10.69767 |
| Japan | 40 | Q2 | 6 | 215 | 2.790698 |
| Japan | 40 | Q3 | 21 | 215 | 9.767442 |
| Japan | 40 | Q4 | 100 | 215 | 46.51163 |
| Japan | 40 | Q5 | 7 | 215 | 3.255814 |
| Japan | 40 | Q6 | 22 | 215 | 10.23256 |
| Japan | 80 | Change OCC | 8 | 215 | 3.72093 |
| Japan | 80 | Change JC | 10 | 215 | 4.651163 |
| Japan | 80 | Neg. Infinite | 6 | 215 | 2.790698 |
| Japan | 80 | No change | 6 | 215 | 2.790698 |
| Japan | 80 | Pos. Infinite | 13 | 215 | 6.046512 |
| Japan | 80 | Q1 | 35 | 215 | 16.27907 |
| Japan | 80 | Q2 | 4 | 215 | 1.860465 |
| Japan | 80 | Q3 | 22 | 215 | 10.23256 |
| Japan | 80 | Q4 | 95 | 215 | 44.18605 |
| Japan | 80 | Q5 | 7 | 215 | 3.255814 |
| Japan | 80 | Q6 | 9 | 215 | 4.186047 |
| Japan | 160 | Change OCC | 14 | 211 | 6.635071 |
| Japan | 160 | Change JC | 11 | 211 | 5.21327 |
| Japan | 160 | Neg. Infinite | 4 | 211 | 1.895735 |
| Japan | 160 | No change | 14 | 211 | 6.635071 |
| Japan | 160 | Pos. Infinite | 12 | 211 | 5.687204 |
| Japan | 160 | Q1 | 30 | 211 | 14.21801 |
| Japan | 160 | Q2 | 4 | 211 | 1.895735 |
| Japan | 160 | Q3 | 13 | 211 | 6.161137 |
| Japan | 160 | Q4 | 93 | 211 | 44.07583 |
| Japan | 160 | Q5 | 7 | 211 | 3.317536 |
| Japan | 160 | Q6 | 9 | 211 | 4.265403 |

**Table S7. Change in occupancy over time**

| **Study area** | **Grain Size** | **OCC slope** | **SE** | **p-value** | **R-squared** |
| --- | --- | --- | --- | --- | --- |
| Czechia | 11 | -0.77269 | 0.796738 | 0.332502 | 0.001456 |
| Czechia | 22 | -0.28985 | 0.22131 | 0.190759 | 0.002652 |
| Czechia | 44 | -0.06482 | 0.062345 | 0.298854 | 0.001673 |
| Europe | 50 | 1.26568 | 1.296788 | 0.329329 | 0.001086 |
| Europe | 100 | 0.30403 | 0.383108 | 0.42765 | 0.000718 |
| Europe | 200 | 0.05405 | 0.108761 | 0.619346 | 0.000282 |
| Europe | 400 | 0.01317 | 0.047731 | 0.782702 | <0.01 |
| Europe | 800 | 0.00548 | 0.011765 | 0.641244 | 0.000248 |
| New York | 5 | 2.42558 | 7.820981 | 0.756588 | 0.000197 |
| New York | 10 | 0.5297 | 2.350599 | 0.821805 | 0.000104 |
| New York | 20 | 0.04456 | 0.67501 | 0.947396 | <0.01 |
| New York | 40 | -0.02288 | 0.19355 | 0.905932 | <0.01 |
| New York | 80 | -0.00039 | 0.058516 | 0.994686 | <0.01 |
| Japan | 20 | 0.55522 | 1.054725 | 0.598851 | 0.000598 |
| Japan | 40 | 0.27616 | 0.404579 | 0.495206 | 0.001001 |
| Japan | 80 | 0.12571 | 0.145743 | 0.388831 | 0.001597 |
| Japan | 160 | 0.04535 | 0.050783 | 0.372315 | 0.001705 |

**Table S8. Proportion of edge cells in each atlas**

| **Study area** | **Grain Size** | **N Cells** | **N Edge Cells** | **Proportion Edge Cells** |
| --- | --- | --- | --- | --- |
| Czechia | 11 | 628 | 147 | 0.234076 |
| Czechia | 22 | 176 | 68 | 0.386364 |
| Czechia | 44 | 54 | 33 | 0.611111 |
| Europe | 50 | 2821 | 1072 | 0.380007 |
| Europe | 100 | 810 | 424 | 0.523457 |
| Europe | 200 | 231 | 160 | 0.692641 |
| Europe | 400 | 106 | 92 | 0.867925 |
| Europe | 800 | 30 | 29 | 0.966667 |
| New York | 5 | 5319 | 607 | 0.114119 |
| New York | 10 | 1396 | 264 | 0.189112 |
| New York | 20 | 384 | 126 | 0.328125 |
| New York | 40 | 112 | 61 | 0.544643 |
| New York | 80 | 37 | 30 | 0.810811 |
| Japan | 20 | 1184 | 539 | 0.455236 |
| Japan | 40 | 376 | 235 | 0.625 |
| Japan | 80 | 124 | 104 | 0.83871 |
| Japan | 160 | 43 | 43 | 1 |

**Figure S1. Slope of the change in Join count difference (JCD) with time across grain sizes.** Slope values of the regressions of the JCD against time across grain sizes (represented by the length of the side of the cell), each line representing a different region. The x-axis is on a log_10_ scale. Negative slopes indicate a decline in SAC over time, while positive slopes indicate an increase in SAC. The dashed line at zero represents no net change in SAC.

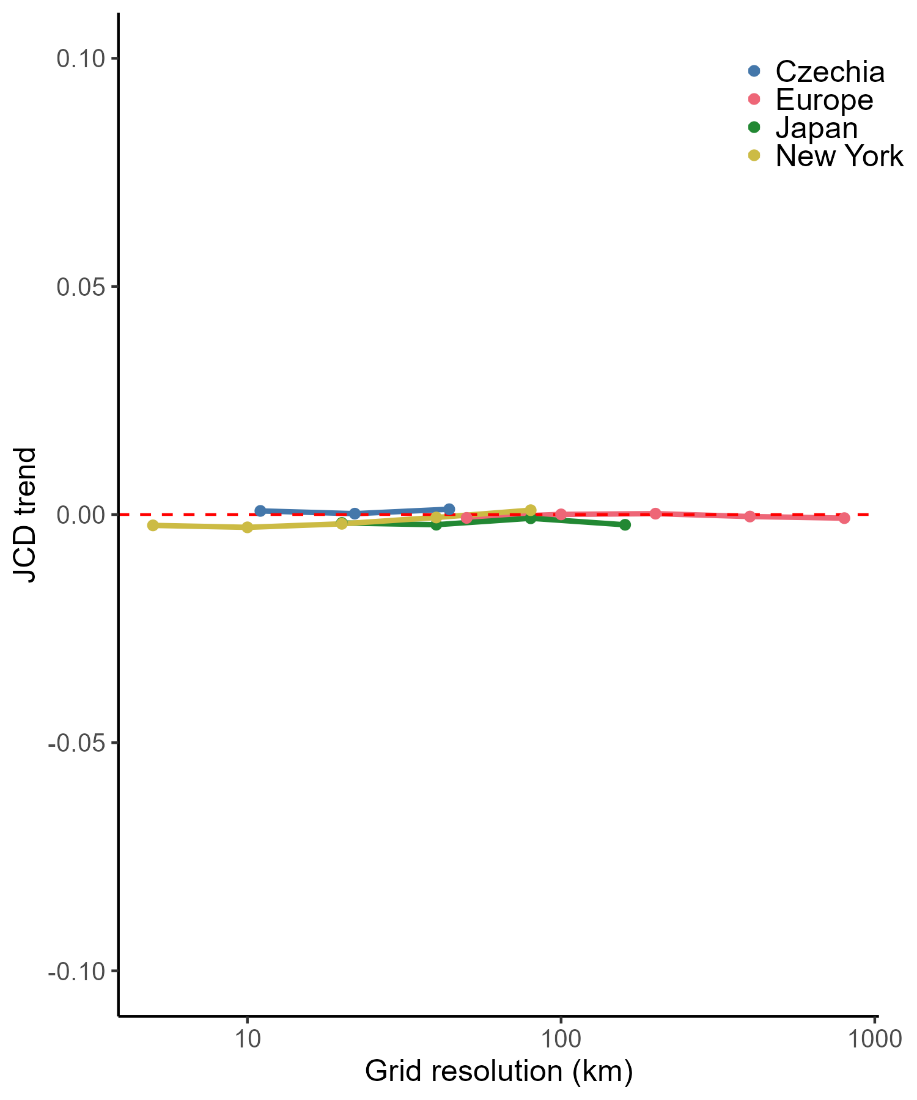

**Figure S2. Median Join count difference (JCD) of species distributions across grain sizes.** Each line represents an atlas location and replication. Line colors indicate atlas location, and symbols and line types represent atlas periods.

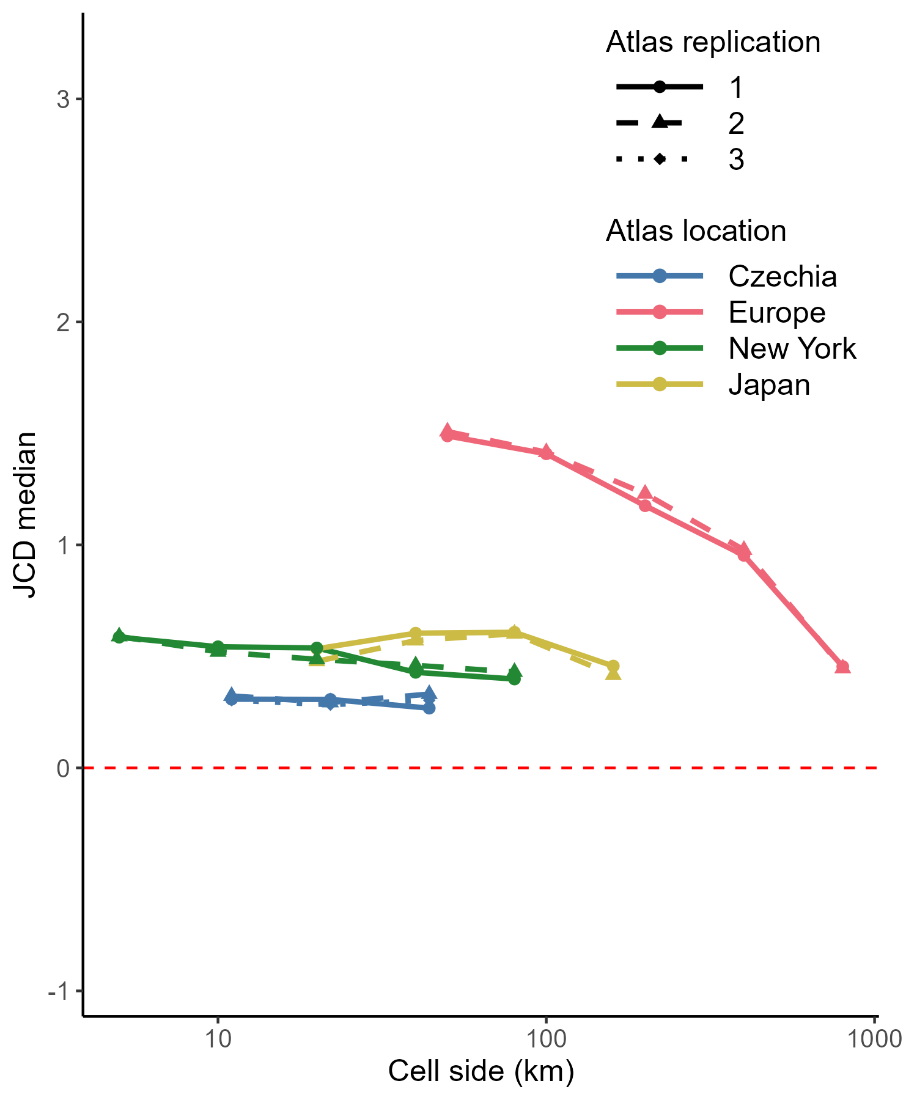

**Figure S3. Percentage of species in each quadrant of joint change in occupancy and JC across grain sizes per atlas location.**
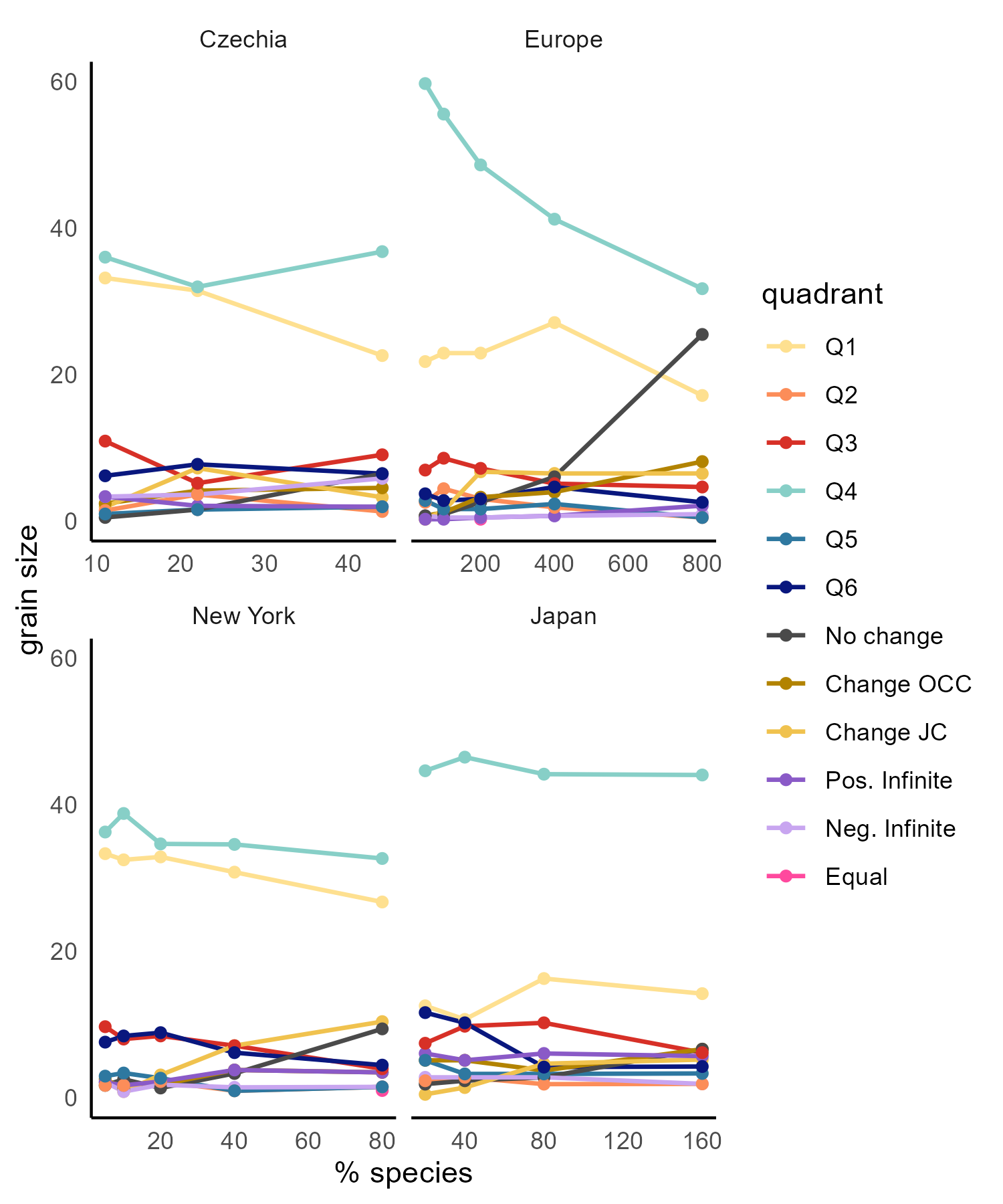
